# Lactation is an Exercise-Responsive Window for Intergenerational Metabolic Programming

**DOI:** 10.64898/2026.11.23.768661

**Authors:** Shiana S. Loona, Vidhi D. Parmar, Katherine A. Klaus, Ameya J. Petsch, Kyle J. Sevits, Robert G. Leija, Shankarappa Manjunatha, Giovanni Barazzoni, Hawley E. Kunz, Mark W. Pataky

## Abstract

Maternal exercise during pregnancy can influence offspring metabolic health, but whether exercise initiated exclusively during lactation has lasting effects is less clear. We investigated the effects of postpartum exercise in chow-fed and high-fat diet (HFD)-fed dams and followed their offspring after weaning. Female C57BL/6J mice consumed chow or a 60% HFD before mating and throughout lactation. Beginning on postpartum day 1, dams were assigned to sedentary or treadmill-exercise conditions until weaning on day 25. Offspring were maintained on chow and assessed longitudinally for body mass, body composition, glucose tolerance, and insulin tolerance through adulthood. Postpartum exercise reduced maternal adiposity and improved maternal glucose tolerance, with effects observed in both chow- and HFD-fed dams. Maternal postpartum exercise was associated with altered offspring growth trajectories and body composition, with effects becoming more apparent with advancing age. Glucose tolerance also improved with maternal exercise in adult male and female offspring, whereas insulin tolerance did not differ significantly among maternal treatment groups. These findings suggest that maternal exercise during lactation is associated with long-term differences in offspring metabolic health that persist long after cessation of the maternal intervention and become increasingly apparent in adulthood. Our findings expand the exercise-responsive window for developmental metabolic programming beyond pregnancy and support lactation as a potentially modifiable period during which maternal physical activity may influence long-term offspring metabolic health.

**ARTICLE HIGHLIGHTS:**

- Although postpartum physical activity is recommended for benefits on maternal health, the influence of exercise in lactating mothers on the long-term metabolic health of the nursed offspring has not been previously tested.
- We assessed whether treadmill exercise training initiated after parturition in mouse dams influenced the long-term metabolic health of their offspring.
- In addition to reducing diet-induced maternal adiposity and glucose intolerance, postpartum maternal exercise training reduced body composition and enhanced glucose tolerance in male and female adult offspring.
- The findings identify lactation as a potential window for maternal exercise to influence offspring metabolism, although the mediating signals remain uncertain.

## INTRODUCTION

During pregnancy, the maternal metabolic environment shapes fetal development through placental delivery of nutrients, hormones, and other signaling molecules, thereby influencing susceptibility to chronic disease later in life (1). Among the maternal factors known to influence developmental programming, obesity is one of the most extensively studied. Children born to mothers with obesity exhibit an increased risk of obesity, type 2 diabetes, and cardiovascular disease that persists throughout adulthood (2). Rodent studies have similarly demonstrated that maternal consumption of an obesogenic diet before and during pregnancy results in long-lasting alterations in offspring body composition and glucose homeostasis (3–5). Conversely, maternal exercise during gestation favorably modifies the intrauterine environment and improves offspring metabolic health. Mouse studies have shown that gestational exercise protects offspring from maternal diet-induced glucose intolerance in adulthood (6–8) via direct delivery of placental factors to the growing fetus (9), while human studies associate maternal exercise training during gestation with reduced infant adiposity, greater infant metabolic rate, and reduced odds of macrosomia (10–12). Together, these findings demonstrate that maternal health and behavior during gestation play a critical role in offspring developmental programming, with lasting influence for offspring metabolic health.

The postpartum period also represents a critical developmental window where maternal physiology continues to influence offspring development through lactation. Breast milk is a complex biological fluid that contains not only nutrients but also hormones, metabolites, lipids, immune factors, and numerous other bioactive molecules capable of influencing infant growth, metabolism, immune maturation, and development (13). However, maternal metabolic dysfunction, including obesity and diabetes, alters the molecular composition of breast milk in humans and other mammals (14–16). Furthermore, cross-fostering studies demonstrate that mouse offspring born to lean dams but nursed by obese dams develop impaired metabolic health in adulthood (17; 18), indicating that maternal metabolic health can program offspring metabolism during lactation independently of the intrauterine environment.

Emerging evidence suggests that maternal exercise can modify the lactational environment. Initial evidence from cross-fostering studies showed that mouse offspring nursed by dams that were exercise trained before parturition exhibited improved metabolic health in adulthood, implicating milk-mediated mechanisms of maternal exercise (19). More recently, exercise initiated exclusively during lactation was shown to remodel mammary epithelial metabolism, reverse obesity-associated alterations in milk composition, and enhance neonatal mouse lipid oxidation (20). Likewise, human studies have shown that maternal exercise during lactation alters the breast milk metabolome and other bioactive factors potentially capable of transmitting beneficial molecular cues to the growing infant (21–24) without any negative impact on breast milk production (25; 26), infant acceptance of milk (27), or important immunoregulatory factors in milk (28). Together, these findings suggest that lactation is an exercise-responsive biological window and raise the possibility that maternal postpartum exercise may beneficially modify offspring metabolic developmental programming through alterations in breast milk.

Although gestation is widely recognized as a critical period for developmental programming, it remains unclear whether developmental trajectories can be modified after birth. Specifically, whether maternal exercise initiated exclusively during lactation is sufficient to reshape offspring metabolic programming despite prior exposure to an obesogenic intrauterine environment remains unknown. Resolving this question would establish whether lactation represents not only a developmental window, but also a therapeutic window during which maternal behavior can improve long-term offspring metabolic health. We therefore investigated the effects of maternal exercise performed exclusively during lactation on metabolic health in adult offspring born to chow-fed and high-fat-fed dams.

## RESEARCH DESIGN AND METHODS

### Animal Management

Mayo Clinic Institutional Animal Care and Use Committee approved this research. At 6 weeks of age, female C57BL/6J mice were randomized to standard chow or a 60% HFD. After 2 weeks, females were mated with age-matched, chow-fed males and remained on their assigned diets throughout mating, gestation, and lactation (Fig. 1A). Following parturition, litters were culled to 7 pups to ensure adequate access to milk, and dams within each dietary group were randomized to either a sedentary (Sed) or exercise-trained (Ex) condition, generating four groups: Sed-HFD, Ex-HFD, Sed-Chow, and Ex-Chow, (n = 7-9 dams/group). The postpartum exercise intervention began one day postpartum and continued until weaning on postpartum day 25. Exercise-trained dams completed a progressive treadmill-running program twice daily, 5 days/week throughout lactation (Supplementary Fig. 1A). Each exercise session consisted of a brief warm-up (beginning at 10 m/min) followed by progressively increasing running speed at constant grade. Two daily sessions were used to limit prolonged dam-pup separation, one in the morning (~7:00-8:00) and one in the afternoon (~16:00-17:00), separated by approximately 9 hours. Duration, speed, and treadmill grade were progressively increased for the first 14 days, and then maintained at a consistent duration, speed, and grade until weaning. Immediately following each exercise session, dams were returned to their litters. Sedentary dams were separated from their litters and placed in empty individual cages (without bedding or water) for a duration matched to the exercise sessions to control for the potential effects of maternal-pup separation and handling experienced by the exercising dams. Dam body mass and food consumption were measured at regular intervals and were performed in the morning to minimize circadian variation in these outcomes. On postpartum day 25, offspring were weaned, and dam body composition was assessed. Dams were then fasted overnight before glucose tolerance testing on the following morning (postpartum day 26). Within 1-2 days (postpartum days 27-28), dams were fasted for 4 h (6:00-10:00) and anesthetized with pentobarbital (50-80 mg/kg, intraperitoneal). Once a surgical plane of anesthesia was confirmed, exsanguination was performed, and parametrial white adipose tissue was carefully excised, weighed, and processed for adipocyte size analysis. Following tissue collection, the heart was excised to complete euthanasia.

**Figure 1:**
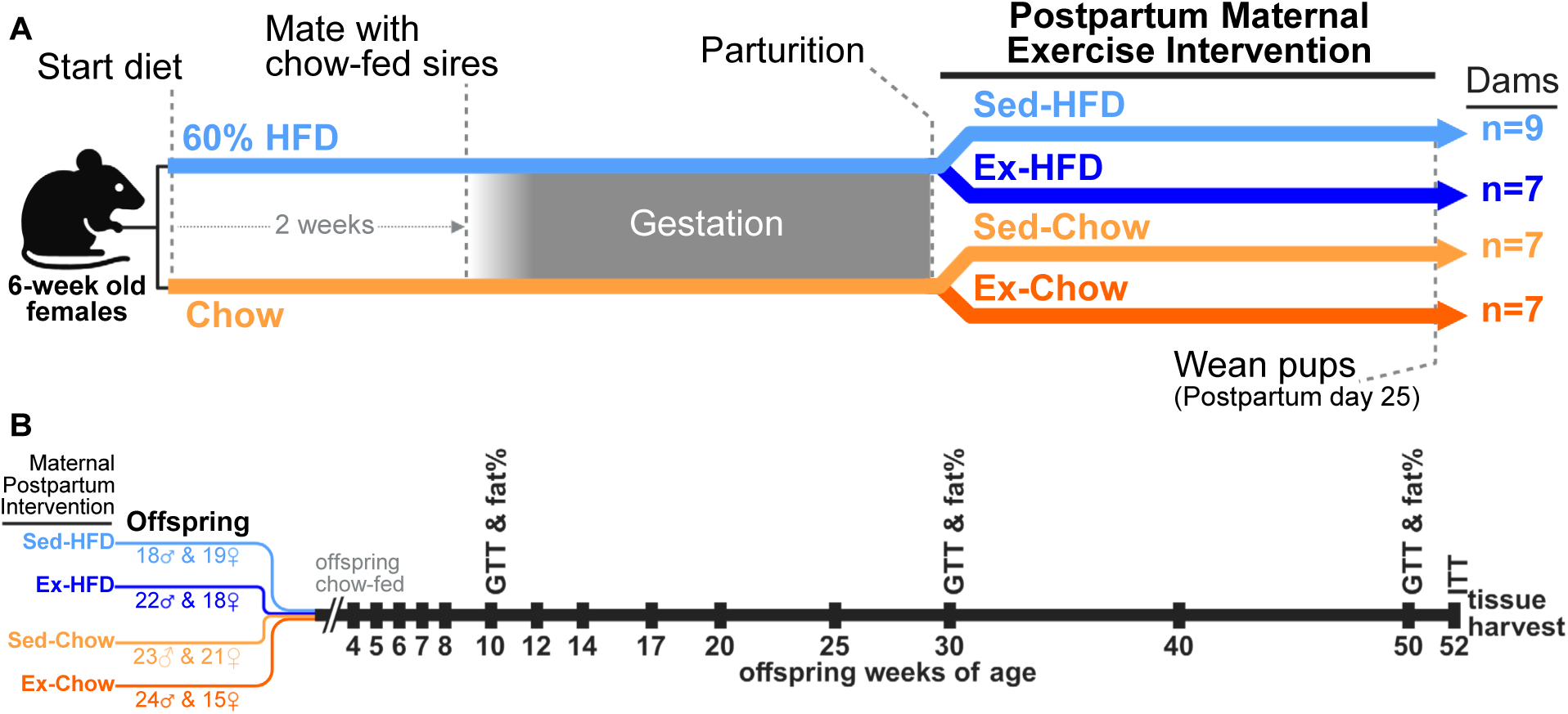
Study Design. (**A**) Non-pregnant female C57BL/6J mice were fed a chow or high-fat diet (HFD) for two weeks prior to and during mating with chow-fed sires. After parturition, dams were maintained on their respective diets and randomized into either sedentary (Sed) or postpartum exercise training (Ex), resulting in four treatment groups Sed-HFD, Ex-HFD, Sed-Chow, and Ex-Chow (n=7-9 dams per group). Created in Biorender. (**B**) Following weaning (25 days postpartum), all offspring were maintained on a chow diet. Male and female offspring from Sed-HFD (18 male, 19 female), Ex-HFD (22 male, 18 female), Sed-Chow (23 male, 21 female), and Ex-Chow (24 male, 15 female) dams were monitored longitudinally until 1 year of age. Intraperitoneal glucose and insulin tolerance tests (GTT and ITT) as well as body composition (fat%) were measured at the indicated weeks. Body mass was measured at each of the weeks with number label. Created in Biorender.

Weaned offspring were group-housed by sex (<5 mice/cage). Offspring from different maternal treatment groups were co-housed when birth dates were within 1 day. Mice were individually identified within each cage using unique ear-clip patterns. Regardless of maternal treatment group, all offspring were maintained on a standard chow diet for the duration of the study. Male and female offspring were statistically analyzed separately, and all weaned offspring were included in the study. Because multiple offspring originated from the same dam, dam (litter) was included as a random effect in linear mixed-effects models to account for within-litter correlation and unequal numbers of same-sex offspring per litter (see Statistical Analysis below). Body mass, body composition, and glucose tolerance were measured longitudinally (Fig. 1B). Offspring insulin tolerance weas measured at 52 weeks of age. Within 2-7 days after the insulin tolerance test, offspring underwent a 4-hour fast (6:00-10:00) before terminal tissue collection. Mice were anesthetized via intraperitoneal (i.p.) injection of pentobarbital (50-80 mg/kg), and exsanguination was performed by cardiac puncture prior to excision of the heart. Livers were rapidly dissected and mitochondria were isolated and assessed. Perigonadal adipose tissue was then rapidly dissected and processed for subsequent analyses.

### Metabolic Phenotyping

For all tolerance tests, mice had ad libitum access to water during fasting. Glucose tolerance tests (GTT) were conducted in dams and offspring (at 10, 30 and 50 weeks of age) after an 11-hour overnight fast (20:00-7:00). Tail blood samples for glucose measurements from the GTT were obtained before and 15, 30, 60, and 120 minutes after i.p. glucose injection (2 g/kg body mass). Insulin tolerance tests (ITT) were conducted in offspring (at 52 weeks of age) by i.p. insulin injection (1.5 U/kg fat free mass) after a 2-hour fast (8:00-10:00). Tail blood samples for glucose measurements from the ITT were obtained before and 10, 20, 30, 45, 60, 90, and 120 minutes after i.p. insulin injection. Blood glucose levels were measured using an AlphaTrak 2 glucometer (Abbott Laboratories). Area under the curve (AUC) was calculated to evaluate glucose and insulin tolerance.

### Body Composition

Body composition was measured non-invasively by nuclear magnetic resonance using a Burker Minispec LF90 II (Bruker, Billerica, MA, USA). Mice were placed individually in a restraining tube without anesthesia, and analyzed according to the manufacturer’s instructions.

### Mitochondria Isolation

Mitochondria were isolated from approximately 200mg liver tissue. Liver tissue was weighed and transferred into 1 ml of ice-cold Liver Buffer A consisting of 220mM mannitol, 70mM sucrose, 0.5mM EGTA, 2 mM HEPES and 0.1% fatty acid free BSA (pH 7.2). The tissue was then minced and diluted into 6ml total volume and homogenized with a Teflon/glass homogenizer for 2 min on ice water. Homogenate was then centrifuged at 720g for 5 min. The supernatant was collected in a new tube and centrifuged at 9000g for 5 min. The resulting pellet was suspended and rinsed in 1 ml of Liver Buffer A and again centrifuged at 9000 g for 5 min. The resultant pellet was resuspended in Buffer B (180 mM sucrose, 35mM KH_2_PO_4_, 10mM magnesium acetate, 5mM EDTA, pH 7.2) with a dilution factor of initial tissue weight x 4.2.

### Mitochondrial oxygen consumption

Mitochondrial respiration was measured using Oxygraph-O2K-Fluorescence LED2-Module (Oroboros Instruments, Innsbruck, Austria), as previously described (22; 29). Oxygen consumption rate (OCR) was measured in a 50 μl aliquot of isolated mitochondria suspension added to the 2 ml Oxygraph chamber and allowed to equilibrate. Respiration was measured devoid of substrates (state 1; mitos); in the presence of 10 mM glutamate, 2 mM malate, and 10 mM succinate (state 2; G+M+S); and 2.5 mM ADP (state 3, complex I and II). 0.5µM rotenone was added, followed by addition of 2 μg/ml oligomycin. Uncoupling was measured using 0.01mM titrations of FCCP. Finally, 2.5 μM antimycin A was added. OCR was normalized per microgram of mitochondrial protein, as determined using the DC Protein Assay (Bio-Rad Laboratories, Hercules, CA, USA).

### Mitochondrial ATP production

Mitochondrial ATP production was measured enzymatically by coupling ATP formation to NADPH production through hexokinase and glucose-6-phosphate dehydrogenase, as previously described (30). NADPH autofluorescence was continuously measured using a Fluorolog 3 spectrofluorometer (Horiba Scientific, Piscataway, NJ, USA). Isolated mitochondrial suspension (10 µl) was added to 2 ml buffer Z containing 110 mM K-MES, 35 mM KCl, 1 mM EGTA, 5 mM K_2_HPO_4_, 3 mM MgCl_2_-6H_2_O, and 5 mg/ml bovine serum albumin (pH 7.4, 295 mOsm) and 2.5 mM D-Glucose. State 3 ATP production was normalized to mitochondrial protein content.

### Adipocyte Cell Size

Adipocyte size was measured as previously described (31). Briefly, 1 mg/mL collagenase type II (Sigma, C-6885) was added to ~100 mg of fresh perigonadal adipose tissue in HEPES buffer (0.1 M HEPES, 0.12 M NaCl, 0.05 M KCl, 0.005 M glucose, 1.5% w/v BSA, 1 mM CaCl2, pH 7.4). Adipose was digested in a 37°C water bath for 20 min, centrifuged (300 g, 2 min), and a 50–150 µL aliquot of the resultant floating adipocyte layer was incubated with 450 µL of 0.2% methylene blue in HEPES buffer at 37°C for 5 min. Stained adipocytes were mounted on glass slides and imaged at ×10 magnification. Adipocyte area, diameter, and volume were quantified using the Cell Counting and Analysis Program (Mayo Clinic Biomedical Imaging Resource). Mean adipocyte lipid content was calculated from >300 adipocytes/sample as adipocyte volume × 0.915 (density of triolein) (31; 32). For two samples, 218 (male, Chow-Sed) and 240 (male, Chow-Ex) cells were quantified because fewer images were available. Their mean cell sizes were not outliers and were retained in the analyses.

### Statistical Analyses

Data are presented as means with 95% confidence intervals (CIs), and statistical significance was defined as P < 0.05. Maternal and litter-level outcomes (litter size at birth and number of pups surviving to weaning) were analyzed using two-way analysis of variance (ANOVA), with maternal diet (chow or HFD), postpartum exercise (Sed or Ex), and their interaction as fixed effects. Offspring outcomes were analyzed separately by sex using linear mixed-effects models (LMMs), with maternal diet, postpartum exercise, and their interaction included as fixed effects and dam (litter) as a random effect (33). Because multiple offspring originated from each dam, individual offspring measurements were treated as clustered within litters rather than as independent replicates of the maternal intervention. This approach accounted for within-litter correlation and accommodated unequal numbers of same-sex offspring per litter while retaining individual offspring observations. Statistical analyses were performed using IBM SPSS Statistics, version 29.0.2.0 (IBM Corp., Armonk, NY, USA).

### Data and Resource Availability

Source data for the graphs are provided in Supplementary Data File S1 containing the values that were used to create all the graphs in the paper. Any additional information required to reanalyze the data reported in this paper is available from the lead contact upon request. This study did not generate any unique reagents.

## RESULTS

### Postpartum Exercise Training Mitigates Glucose Intolerance and Excessive Adipose Retention in Obese Lactating Dams

Dams that were fed a HFD had greater body mass than chow-fed dams, and exercise training mitigated the postpartum gain in body mass (Fig. 2A). Despite these differences in body mass, food intake increased during lactation and was generally similar among groups (Fig. 2B). At weaning, maternal body fat percentage was significantly influenced by diet (main effect of diet; P<0.001) and reduced by postpartum exercise (main effect of exercise; P=0.005), with a trend (P=0.063) towards a significant diet-by-exercise interaction (Fig. 2C). Parametrial fat mass was significantly lower in chow-fed dams (Supplementary Fig. S1B), and parametrial adipocyte volume was significantly lower in chow-fed versus HFD-fed (P < 0.001), and in exercise trained versus sedentary (P = 0.014) dams (Fig. 2D). HFD feeding impaired dam glucose tolerance, but postpartum exercise training greatly mitigated the diet-induced glucose intolerance in HFD-fed dams (Fig. 2E-F). Thus, exercise initiated immediately after parturition attenuated several adverse maternal metabolic outcomes by the end of lactation.

**Figure 2:**
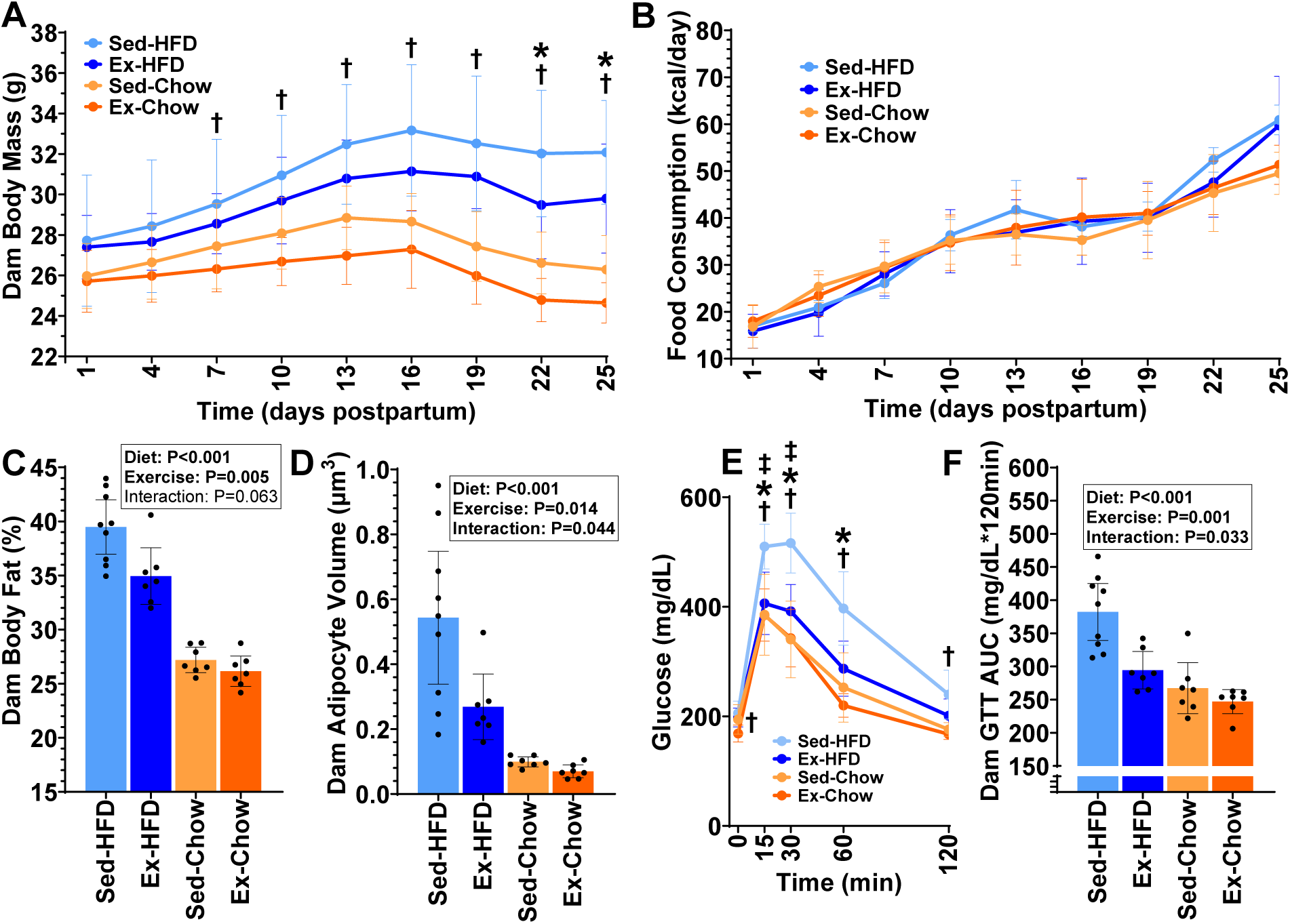
Postpartum Exercise Training Mitigates Glucose Intolerance and Excessive Adipose Retention in Obese Lactating Dams. (**A-B**) Body mass and food consumption measured every 3 days during the postpartum intervention. (**C**) Dam body composition measured on the day of weaning (25 days postpartum). (**D**) Dam mean adipocyte volume determined from the parametrial fat pad (dissected at 2-3 days post weaning). (**E-F**) Dam blood glucose (E) following intraperitoneal glucose injection (2 g/kg body mass) performed after an 11 hour overnight fast. Glucose area under the curve (AUC) is presented (F). All data were analyzed by 2-way ANOVA. P-values for main effects of diet, exercise, and diet × exercise interaction are shown in panels C, D, and F. For panels A and E, * indicates a significant (P<0.05) main effect of exercise, † indicates a significant (P<0.05) main effect of diet, and ‡ indicates a significant (P<0.05) the diet × exercise interaction at each timepoint. All data are presented as mean ± 95% CI, and dots are presented to indicate individual dams when appropriate.

### Litter outcomes

Litter size (number of pups born per litter) did not differ significantly by maternal diet (P = 0.858; Supplementary Fig. S1C). Litters with more than seven pups were culled to seven within 1 day after parturition (Supplementary Fig. S1D). The number of pups surviving to weaning was significantly lower in litters nursed by HFD-fed dams (P = 0.046), but pup survival was improved if HFD-fed dams were postpartum exercise trained, as indicated by a significant diet-by-exercise interaction (P = 0.024; Supplementary Fig. S1D).

### Postpartum Maternal Exercise Training Improves Offspring Growth Trajectory and Body Composition Throughout the Lifespan

In both male (Fig. 3A) and female (Fig. 3B) offspring, maternal postpartum exercise during lactation resulted in significantly lower body mass by 5 weeks of age, regardless of maternal diet, and this effect persisted throughout the offspring lifespan. In male offspring, but not female offspring, maternal HFD reduced body mass through 20 weeks of age compared to male offspring of chow-fed dams (Fig. 3A).

**Figure 3:**
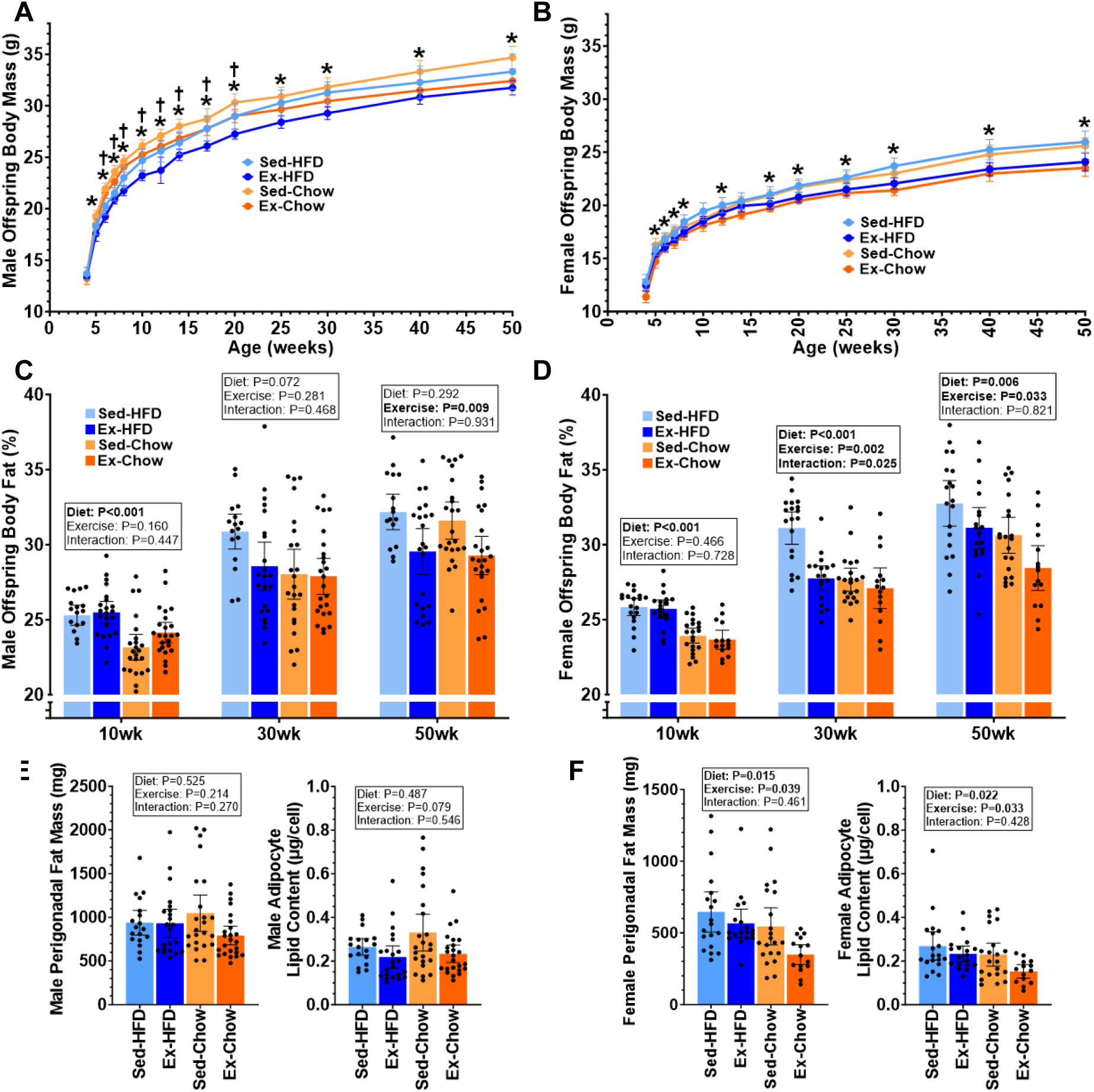
Postpartum Maternal Exercise Training Improves Offspring Growth Trajectory and Body Composition Throughout the Lifespan. (**A-B**) Male (A) and female (B) offspring body weight measured longitudinally from weaning through 50 weeks of age. (**C-D**) Male (C) and female (D) offspring body fat percentage measured at 10, 30, and 50 weeks of age. (**E-F**) Male (E) and female (F) offspring perigonadal fat mass (left panel) and adipocyte lipid content (right panel) measured at sacrifice (52 weeks of age). All offspring data were analyzed using linear mixed-effects models with dam diet, dam postpartum exercise, and the diet × exercise interaction included as fixed effects and litter included as a random effect to account for clustering of littermates. In panels A-B, * indicates a significant (P<0.05) main effect of exercise and † indicates a significant (P<0.05) main effect of diet at each timepoint. In panels C-F, P-values for the main effects of diet, exercise, and the diet × exercise interaction are shown. All data are presented as mean ± 95% CI. In panels C-F, dots represent individual offspring.

In both male (Fig. 3C) and female (Fig. 3D) offspring of HFD-fed dams, body-fat percentage at 10 weeks of age was significantly greater than offspring of chow-fed dams (P < 0.001). The effect of maternal HFD on male offspring body composition was absent by 30 weeks of age, but persisted in female offspring through 50 weeks of age. Interestingly, although maternal postpartum exercise had no effect on either male or female offspring body composition at 10 weeks of age, there was a main effect of maternal postpartum exercise to significantly reduce male offspring body fat percentage at 50 weeks of age, and female offspring body fat percentage at 30 and 50 weeks of age. When offspring were sacrificed at 52 weeks of age, male offspring perigonadal (epididymal) fat mass and adipocyte lipid content were not significantly affected by maternal diet or exercise, although there was a trend (P = 0.079) towards a significant effect of maternal exercise for lower adipocyte lipid content (Fig. 3E). On the other hand, there were significant main effects of both maternal diet and exercise in 52-week-old female offspring for reduced perigonadal (parametrial) fat mass and adipocyte lipid content (Fig. 3F).

### Postpartum Maternal Exercise Training Improves Adult Offspring Glucose Tolerance

In both male and female 10-week-old offspring, glucose tolerance did not differ significantly based on maternal diet or postpartum exercise training (Figs. 4A-D). At 30 weeks of age, glucose tolerance was significantly improved in offspring of chow-fed compared to HFD-fed dams (main effect of diet: males, P = 0.033; females, P = 0.014). Maternal postpartum exercise training also significantly improved glucose tolerance in 30-week-old male offspring (main effect of exercise, P = 0.040), with a similar trend in female offspring (P = 0.080). At 50 weeks of age, maternal exercise improved glucose tolerance in both male and female offspring (main effect of exercise: males, P = 0.033; females, P = 0.039), and maternal HFD versus chow diet significantly impaired 50-week-old female offspring glucose tolerance (main effect of diet, P = 0.015), with a similar trend in 50-week-old male offspring (P = 0.094). Whereas adult offspring glucos tolerance was substantially impacted by maternal behaviors (diet and exercise) before and during lactation, insulin tolerance was unaffected by maternal diet or exercise in 52-week-old male and female offspring (Supplementary Figs. 2A-B).

**Figure 4:**
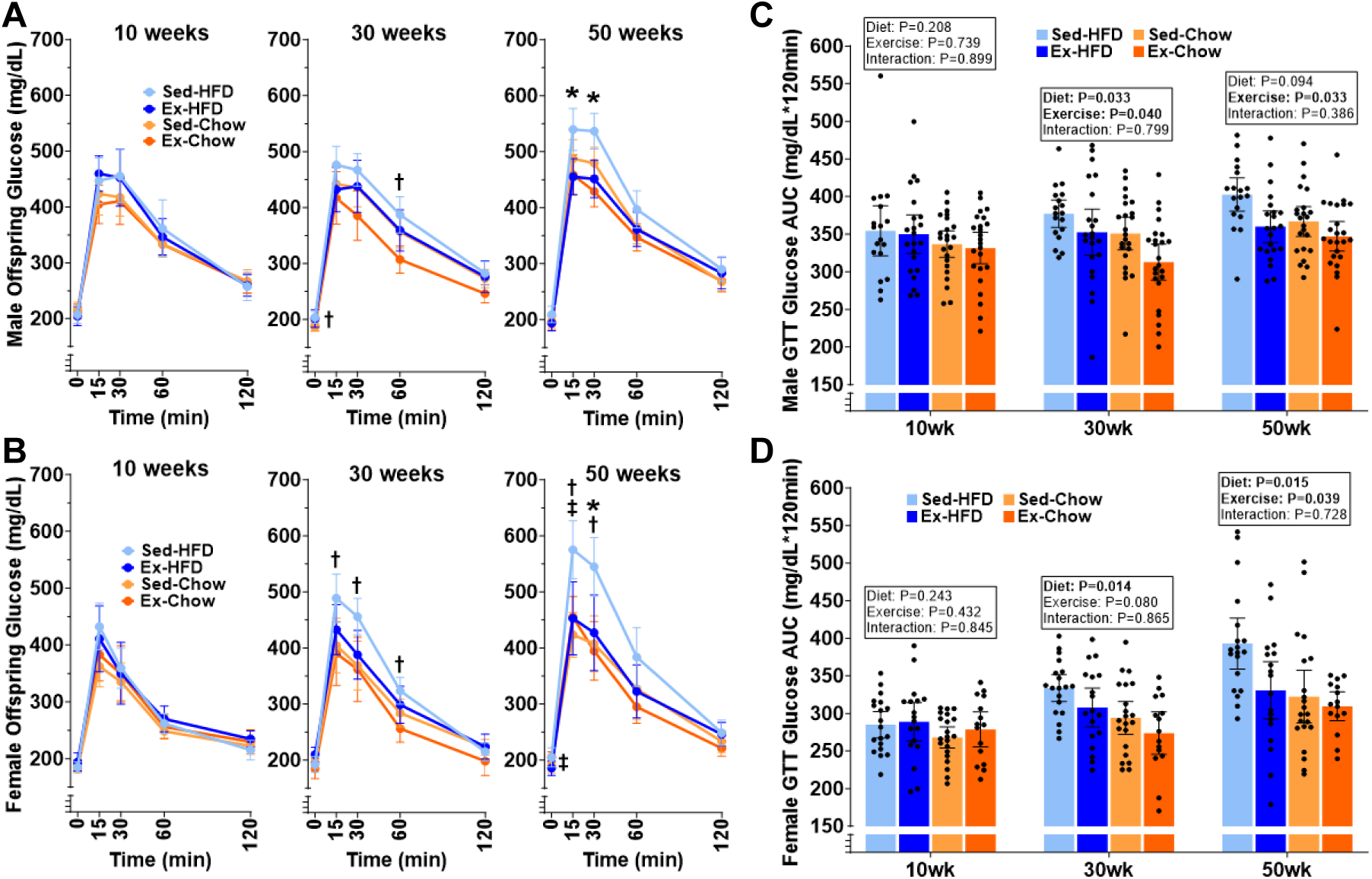
Postpartum Maternal Exercise Training Improves Adult Offspring Glucose Tolerance. (**A-B**) Blood glucose concentrations following intraperitoneal glucose injection (2 g/kg body mass) performed after an 11-hour overnight fast in male (A) and female (B) offspring at 10, 30, and 50 weeks of age. (**C-D**) Blood glucose area under the curve (AUC) from panels A-B are displayed for male (C) and female (**D**) offspring at 10, 30, and 50 weeks of age. All offspring data were analyzed using linear mixed-effects models with dam diet, dam postpartum exercise, and the diet × exercise interaction included as fixed effects and litter included as a random effect to account for clustering of littermates. In panels A-B, * indicates a significant (P<0.05) main effect of exercise, † indicates a significant (P<0.05) main effect of diet, and ‡ indicates a significant (P<0.05) the diet × exercise interaction at each timepoint. In panels C-D, P-values for the main effects of diet, exercise, and the diet × exercise interaction are shown. All data are presented as mean ± 95% CI. In panels C-D, dots represent individual offspring.

We assessed hepatic mitochondrial respiration and ATP production in male and female offspring at approximately 52 weeks of age (Supplementary Figs. 3A-D). Neither outcome showed significant main effects of maternal diet or exercise in either sex. However, diet-by-exercise interactions were detected in females, but not males, for respiration following rotenone (P = 0.011), oligomycin (P < 0.001), and FCCP (P = 0.011), and for ATP production rate (P = 0.006). For these outcomes, mean rates were higher in offspring of exercise-trained versus sedentary chow-fed dams, whereas the opposite pattern occurred in offspring of HFD-fed dams.

## DISCUSSION

The primary finding from the present study was that postpartum maternal exercise can influence offspring metabolic outcomes long after weaning and into adulthood. Exercise training during the postpartum period in the lactating dam reduced maternal adiposity and improved maternal glucose tolerance, while offspring of both sexes exhibited lower adiposity and improved glucose tolerance in adulthood. These findings extend the potential window for maternal exercise-mediated metabolic programming beyond pregnancy and support lactation as an interventional window during which maternal exercise can shape long-term offspring metabolic health.

Maternal HFD in our study increased adult offspring adiposity and impaired glucose tolerance, whereas postpartum exercise attenuated several of these abnormalities. Consistent with our finding, diet-induced maternal obesity has been demonstrated to increase maternal adiposity, hyperglycemia, and milk leptin, accompanied by offspring adipocyte hypertrophy and progressive metabolic dysfunction, despite offspring chow feeding after weaning (3). Thus, the lactational environment can shape offspring metabolic phenotype (34). These findings raise the possibility that postpartum exercise may influence offspring development by improving maternal metabolic health and modifying exercise-responsive signals during lactation.

Previous studies have demonstrated that maternal exercise during gestation can improve adult offspring glucose tolerance, adiposity, and tissue-specific metabolic function, including under conditions of maternal HFD exposure (6; 7; 9; 35). More recently it was demonstrated that exercise initiated during lactation induces mammary metabolic remodeling, alters milk fatty acid composition, and enhances neonatal lipid oxidation (20). Our study extends these lactation-focused findings by demonstrating improvements in adult offspring metabolic health, suggesting that developmental programming for lifelong metabolic health is affected by maternal physical activity behavior during lactation. Because exercise was initiated only after parturition, gestational exercise was not the cause for the observed outcomes. Whereas gestational exercise can influence fetal development through exercise-responsive maternal and placental signals (9; 35), the intervention studied here began after parturition, excluding direct fetal exposure to exercise during gestation, supporting the concept that maternal exercise during lactation can transmit developmental signals after birth.

An important feature of the study was that the effects of maternal postpartum exercise on offspring outcomes became more apparent with age. Stanford et al. similarly demonstrated that consequences of gestational maternal exercise can evolve across the lifespan, with differences in male offspring health becoming evident at 52 weeks (6). Our findings extend the concept of maternal exercise-mediated intergenerational signaling to the postpartum period in the lactating dam and suggest that its effects may extend beyond transient changes in early postnatal growth. Although the signals through which maternal exercise during lactation influences offspring remain unknown, these data imply breast milk as a plausible mechanism. Breast milk contains nutrients and bioactive factors that support postnatal development, and its composition can vary with maternal metabolic status and physical activity (19–21; 23; 36; 37). Notably, Harris et al. identified 3′-sialyllactose as a milk-derived mediator of beneficial metabolic and cardiac effects in mouse offspring that were nursed by dams that had been exercise trained before and during gestation (19). Although they were not exercising in the postpartum period, these data provide evidence that improved maternal metabolic health (via exercise) contributes to beneficial maternal signals that are transmitted through breast milk. Recently, we demonstrated in humans that postpartum exercise can increase breast milk small extracellular vesicle concentration and their bioactive molecular cargo (22). Thus, there is a precedent for maternal exercise to mediate molecular changes in breast milk that could influence offspring development. However, the current study does not definitively establish milk-signaling as the mediator of the observed offspring phenotype. Changes in maternal metabolism, milk composition or availability, nursing behavior, and maternal care could all contribute to the observed results.

We examined hepatic metabolism and adipose tissue as potential contributors to the long-term offspring metabolic phenotype. The liver participates in the first-pass metabolism of nutrients absorbed from the intestine and delivered through the portal circulation, making it a plausible target of changes in milk-derived nutrient availability and signaling during lactation. Its relevance to maternal exercise-mediated programming is supported by further studies demonstrating improved offspring hepatic glucose metabolism and insulin responsiveness following maternal exercise (7), as well as fetal hepatic adaptations mediated by placental SOD3 (9). Adipose tissue was examined because of its relevance to offspring obesity susceptibility and evidence that maternal diet-induced obesity promotes offspring adiposity and adipocyte hypertrophy, whereas maternal exercise before and during gestation can attenuate offspring fat accumulation (3; 7). Because substantial adipose tissue development in rodents occurs postnatally, adipose development may be particularly relevant to an intervention initiated during lactation. In the present study, maternal diet modified the association between postpartum exercise and hepatic mitochondrial respiration and ATP production in female offspring, whereas corresponding interactions were not detected in males. The opposing patterns across maternal diets suggest context-dependent mitochondrial adaptations rather than a uniform enhancement of mitochondrial capacity. However, their contribution to the improvements in offspring glucose tolerance remains uncertain. Maternal exercise was associated with differences in adipose outcomes in female offspring, although several outcomes were unchanged in males, suggesting key sex-specific differences in the etiology of maternal exercise-mediated metabolic programming. Together, these findings identify hepatic metabolism and adipose development as potential contributors to the offspring phenotype, although tissue-specific metabolic studies will be required to establish their causal roles.

### Limitations

Several limitations warrant consideration. Fewer pups survived to weaning in HFD-Sed litters, potentially reflecting reduced milk availability, maternal nursing behavior, or other developmental influences. Lower early postweaning body mass in male offspring of HFD-fed dams also raises questions about nutrient delivery but does not establish undernutrition, particularly given their greater relative adiposity. Differential survival and nursing load may have influenced subsequent offspring outcomes. Glucose tolerance test dosing by total body mass may have produced different loads relative to lean mass, given the group differences in body composition. Nevertheless, since maternal exercise produced lasting differences in offspring adiposity and glycemic responses, benefits mediated through altered body composition remain biologically meaningful. Furthermore, GTT and ITT findings do not establish tissue-specific insulin sensitivity. Forced treadmill exercise may impose stress unlike voluntary human activity. Duration-matched separation of sedentary dams controlled for time away from pups but not necessarily exercise-related stress. Favorable maternal and offspring metabolic outcomes argue against a net adverse effect on measured endpoints, although stress-related contributions cannot be excluded. Finally, milk-derived mechanisms remain unproven, prenatal and lactational dietary effects cannot be separated, terminal tissue measurements cannot establish developmental timing of the effects, and the findings in chow-fed offspring do not establish protection against subsequent obesogenic exposure or comparable benefits in humans.

### Conclusions

In conclusion, maternal exercise initiated during lactation improved maternal metabolic health and was associated with persistent differences in offspring growth, adiposity, and glucose homeostasis. Several offspring effects became more apparent with age, suggesting that the consequences of postpartum maternal exercise may extend well beyond weaning. These findings identify lactation as a potential window for maternal exercise-mediated influences on long-term offspring metabolic health and provide a basis for further investigation of the maternal signals and tissue-specific mechanisms involved.

## ACKNOWLEDGEMENTS

The authors would like to thank Kelli Lytle for her valuable insights interpreting the results.

## Author Contributions

M.W.P. conceived the study. M.W.P., K.A.K., A.J.P., and K.J.S. performed animal handling. S.S.L., V.D.P., K.A.K., K.J.S., R.G.L., G.B., and H.E.K. processed and analyzed samples. S.S.L., V.D.P., K.A.K., R.G.L., S.M., and M.W.P. performed data analysis and interpreted the data. S.S.L., V.D.P., and M.W.P. drafted the manuscript and all authors contributed to the editing of the manuscript. M.W.P. provided scientific oversight at all stages. M.W.P. provided final editing and approval. M.W.P acquired funding for the experiments.

## Lead contact

Further information and requests for resources and reagents should be directed to and will be fulfilled by the Lead Contact, Mark W. Pataky.

