## Supplementary Figures for "Lactation is an Exercise-Responsive Window for Intergenerational Metabolic Programming"

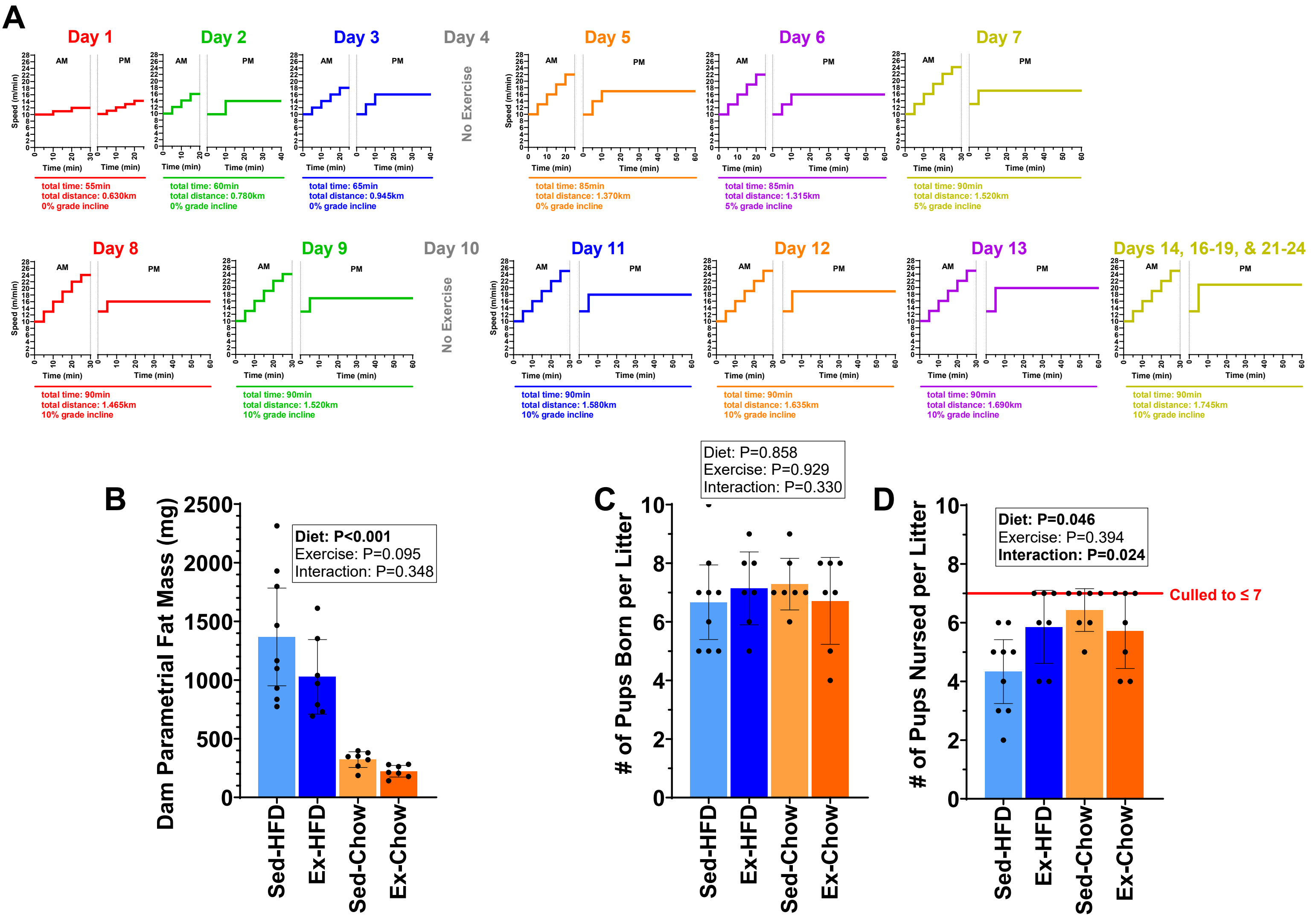


**Supplementary Figure 1: Postpartum Treadmill Exercise Protocol, Maternal Perigonadal Adiposity, and Litter Survival.**

(**A**) Progressive treadmill exercise training protocol performed by postpartum exercise (Ex) dams throughout lactation. Exercise training was initiated on postpartum day 1 and consisted of two daily treadmill running sessions separated by approximately 9 hours on 5 days per week. Running duration and speed were progressively increased over the first 13 days of the intervention before reaching the final training intensity, which was maintained through the remainder of the lactation period. Sedentary (Sed) dams were handled similarly but did not perform treadmill exercise. Morning sessions emphasized exercise intensity, consisting of shorter bouts (20–30 min) at higher running speeds, whereas afternoon sessions emphasized exercise volume, consisting of longer-duration bouts (up to 60 min) performed at a constant moderate running speed. No running was performed on postpartum days 4, 10, 15, and 20 to allow dams to recover from exercise training.

(**B**) Parametrial fat pad mass measured in dams at necropsy (2–3 days following weaning).

(**C**) Litter size at birth.

(**D**) Number of pups surviving to weaning (25 days postpartum). Litters containing more than seven pups were culled to seven pups within 24 hours of birth to normalize litter size across experimental groups.

All data were analyzed by two-way ANOVA. P-values for the main effects of diet, exercise, and the diet × exercise interaction are shown above panels B-D. All data are presented as mean ± 95% CI, and dots represent individual dams.

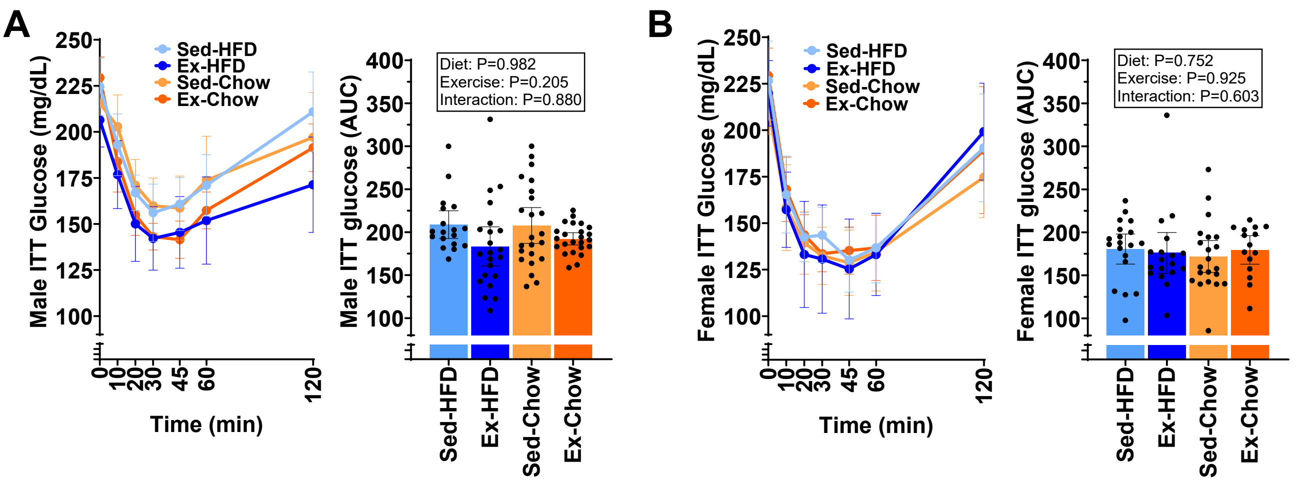


**Supplementary Figure 2: Postpartum Maternal Exercise Training and Adult Offspring Insulin Tolerance.** (**A-B**) Blood glucose concentrations (left panels) following intraperitoneal insulin injection (1.5 U/kg fat free mass) performed after a 2-hour fast in male (A) and female (B) offspring at 52 weeks of age. Blood glucose area under the curve (AUC; right panels) are displayed for male (A) and female (B) offspring at 52 weeks of age.

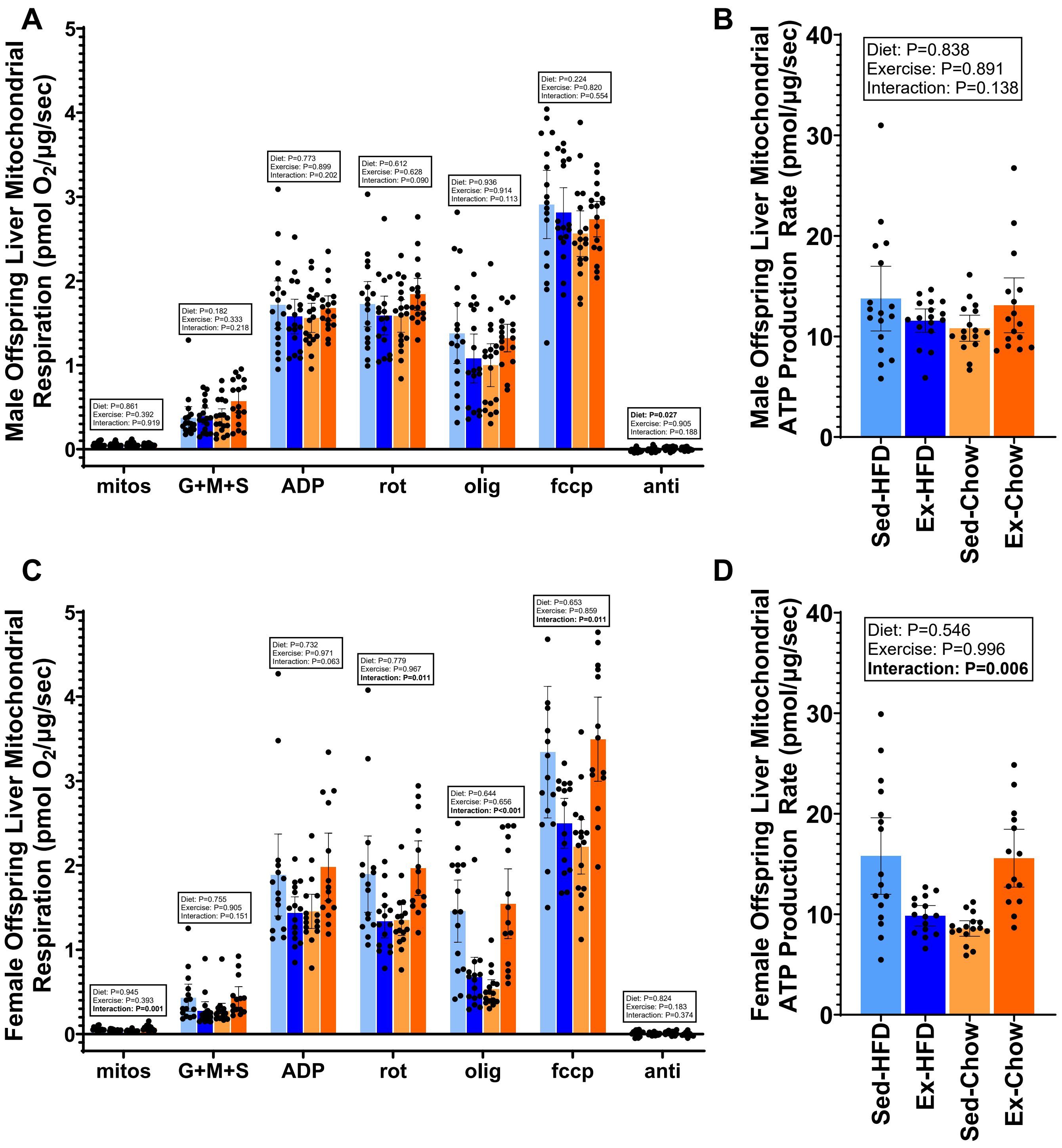


**Supplementary Figure 3: Adult Offspring Hepatic Mitochondrial Respiration and ATP Production Following Maternal Dietary and Postpartum Exercise Interventions.**

(**A-D**) Male (A-B) and female (C-D) offspring oxygen consumption (left panels) and ATP production (right panels) rates measured in isolated liver mitochondria at 52 weeks of age. Respiration is shown before substrate addition (mitos) and following the addition of G+M+S (10 mM glutamate, 2 mM malate, and 10 mM succinate), ADP (2.5 mM), rotenone (rot; 0.5 µM), oligomycin (olig; 2 µg/mL), FCCP (fccp; 0.01 mM titrations), and antimycin A (anti; 2.5 µM). ATP production rate was measured in the presence of ADP (2.5 mM). Oxygen consumption and ATP production rates were normalized to mitochondrial protein content. Male and female offspring were analyzed separately using linear mixed-effects models with maternal diet, maternal postpartum exercise, and the diet × exercise interaction included as fixed effects and litter included as a random effect to account for clustering of littermates. P-values for the main effects of diet and exercise and the diet × exercise interaction are shown above the corresponding measurements. Data are presented as mean ± 95% CI, and dots represent individual offspring.
